# Significant phylogenetic signal is not enough to trust phylogenetic predictions

**DOI:** 10.1101/2022.05.07.490865

**Authors:** Rafael Molina-Venegas, Ignacio Morales-Castilla, Miguel Á. Rodríguez

## Abstract

In a recent study, Cantwell-Jones et al. (2022) proposed a list of 1044 species as promising key sources of B vitamins based primarily on phylogenetic predictions. To identify candidate plants, they fitted lambda models of evolution to edible species with known values in each of six B vitamins (232 to 280 species) and used the estimated parameters to predict B-vitamin profiles of edible plants lacking nutritional data (6460 to 6508 species). The latter species were defined as potential sources of a B vitamin if the predicted vitamin content was ≥15% towards recommended dietary allowances for active females between 31-50 years per 100 g of fresh edible plant material consumed. Unfortunately, the reliability of the predictions that informed the list of candidate species is questionable due to insufficient phylogenetic signal in the data (Pagel’s λ between 0.171 and 0.665) and a high incidence of species with missing values (over 95% of all the species analyzed in the study). We found that of the 1044 species proposed as promising B-vitamin sources, 626 to 993 species showed accuracies that were indistinguishable from those obtained under a white noise model of evolution (i.e. random predictions conducted in absence of any phylogenetic structure) in at least one of the vitamins, which proves the weakness of the inference drawn from imputed information in the original study. We hope this commentary serves as a cautionary note for future phylogenetic imputation exercises to carefully assess whether the data meet the requirements for the predictions to be valuable, or at least more accurate than expected by chance.

## Main

In a recent study, Cantwell-Jones et al.^1^ proposed a list of 1044 species as promising key sources of B vitamins based primarily on phylogenetic predictions. To identify candidate plants, they fitted lambda models of evolution to edible species with known values in each of six B vitamins and used the estimated parameters to predict B-vitamin profiles of edible plants lacking nutritional data. The latter species were defined as ‘sources’ of a B vitamin if the predicted vitamin content was ≥15% towards recommended dietary allowances for active females between 31–50 years per 100 g of fresh edible plant material consumed. This phylogenetic imputation exercise is exciting and inspiring, more so as it would have the potential to be replicated for further goals such as, for example, identifying species of pharmaceutical interest. Unfortunately, the predictions from this study are questionable. The predictive capability of lambda models is primarily determined by the amount of phylogenetic signal in the known data (i.e. the extent to which closely related species share similar values in the trait of interest), which means that predictions based on low phylogenetic signals, even if statistically significant, are typically valueless^2-4^. Further, simulations have shown that even under ‘strong’ phylogenetic signal (i.e. Pagel’s λ = 1), acceptable predictions can only be expected when the most recent common ancestor (MRCA) of a target species and its closest relative with known trait value (hereafter ‘predictive MRCA’) is relatively recent^4^. As such, the older the predictive MRCAs, the lesser the difference between phylogenetic imputations and random predictions (Fig. 1). Based on 232 to 280 nutritionally known species (observed data), Cantwell-Jones et al.^1^ predicted B-vitamin profiles for 6460 to 6508 nutritionally unknown species (over 95% of all the species analyzed in the study), and they did so relying on very weak-to-moderate phylogenetic signals in the observed data (Pagel’s λ between 0.171 and 0.665). These figures suggest that the predictions conducted by the authors are unreliable due to both insufficient phylogenetic signal and a high incidence of relatively old predictive MRCAs (Fig. 1). Moreover, the unreliability of their predictions calls into question the proposed list of 1044 species as promising sources of B vitamins.

**Figure 1.**
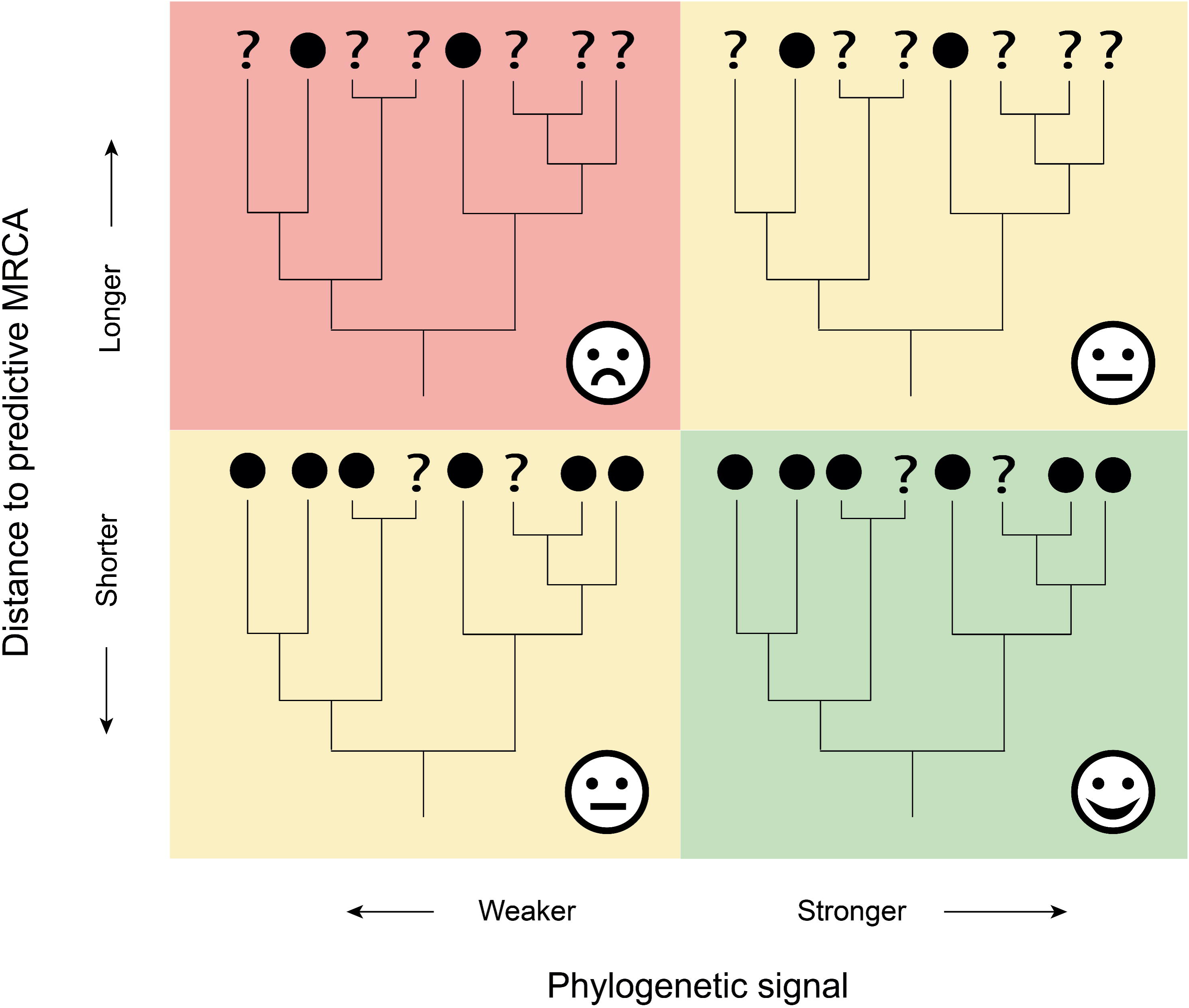
Conceptual model on the expected accuracy of phylogenetic predictions as a function of phylogenetic signal and distance to predictive MRCAs, this is, the MRCA of each target species and its closest relative with known trait value. Accuracy is expected to be acceptable only under strong phylogenetic signal and relatively recent predictive MRCAs (short distances), and predictions will be uncertain if any of these conditions are not met. The probability of finding relatively old predictive MRCAs (long distances) increases with the amount of missing data, which may lead to the worst-case scenario if phylogenetic signal is weak.

Cantwell-Jones et al.^1^ conducted leave-one-species-out cross-validation trials on the observed values of the B vitamins they analyzed (they referred to this procedure as “jackknifing”) and estimated 95% confidence intervals (CI) around each re-estimated (predicted) value. They found that ≥91.4% of nutritionally known species had measured (observed) values within the 95% CI of the values predicted in the trials. Here, we conducted the same analysis but randomizing the authors’ dataset in a two-step procedure. First, the observed values of each vitamin were reshuffled across the species with known value in the vitamins 500 times, and then each reshuffled set (n = 500 sets per vitamin) was checked to show complete lack of phylogenetic signal (i.e. λ < 0.001 and *p* > 0.05). Otherwise, the values were reshuffled iteratively until the conditions were met. We found that ≥92.8% of the observed values in the randomized sets sit within 95% CIs of their corresponding predicted values (Supplementary Data 1-5). This demonstrates that the authors simply found the null expectation of the analysis, which deems the CI-based evidence of their study as invalid.

On the other hand, the authors used generalised least squares models (with a variance structure in the error term) to test the strength of the relationship between predicted and observed values, and they found significant relationships for all the vitamins. Here, we used a simple and more intuitive prediction coefficient (*P*^*2*^) to assess the overall predictive power of the lambda models they employed:

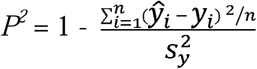

where ŷ_*i*_ and *y*_*i*_ are respectively the predicted and observed value for the nutritionally known species *i* and 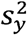 is the sample variance of the observed values of the trait^4-6^. *P*^*2*^ can be interpreted similarly as Ezekiels’ adjusted coefficient of determination^7^, so that *P*^*2*^ = 1 when all predictions perfectly match the observations and *P*^*2*^ = 0 when the model is no better in predicting values than simply taking the mean of the observed values (note that *P*^*2*^ has no negative boundary, as there is no theoretical limit to how badly a model can predict observed trait values). We found that *P*^*2*^ was close to zero in all cases except for folate, which showed a slightly higher (yet weak) score (Table 1). This is not surprising, as previous simulations have shown that phylogenetic predictions perform very poorly when λ is lower than ∼0.6^4^. There are numerous studies that caution against using phylogenies alone to predict missing data under low phylogenetic signal^2-4,8,9^, yet none of them was acknowledged in Cantwell-Jones et al.^1^

**Table 1.** Phylogenetic signal λ of each B vitamin analyzed across *N* nutritionally known species and prediction coefficients *P*^*2*^ derived from the leave-one-species-out cross-validation trials.

| B vitamin | $N$ | $\lambda$ | $P^2$ |
| --- | --- | --- | --- |
| Thiamine | 280 | 0.293 | -0.010 |
| Riboflavin | 277 | 0.192 | 0.035 |
| Niacin | 278 | 0.665 | 0.086 |
| Pantothenic acid | 232 | 0.171 | 0.023 |
| Folate | 256 | 0.302 | 0.183 |

In line with the authors’ observation that “median differences between predicted and observed values for each nutrient were <33% of the standard deviation across species”, it could be argued that some of the predictions may still be useful despite the discouraging results from the leave-one-species-out cross-validation trials (Table 1). Thus, we used the method described in Molina-Venegas et al.^4^ to assess the expected species-level accuracy of the predictions that informed the list of candidate plants (see Supplementary Information for details). Depending on whether *p*-values were Bonferroni-corrected (*n* = 1616, two-sample Wilcoxon tests) and the nominal alpha criterion (5% or 0.1%), we found that of the 1044 species proposed as promising B-vitamin sources, 626 to 993 species showed accuracies that were indistinguishable from those obtained under a ‘white noise’ model of evolution (i.e. random predictions conducted in absence of any phylogenetic structure) in at least one of the vitamins (Supplementary Data 6). Moreover, regarding the species that showed statistically significant accuracies, it should not be concluded that their predictions are necessarily valuable, only that they are better than drawing values at random independently of the phylogeny. This is illustrated by the rather modest maximum accuracy that was recorded across all candidate species and B vitamins analyzed, which corresponded to *Hordeum bulbosum* and niacin with only 54.4% of *P*^*2*^_sim_ values ≥ 0.75 (Supplementary Data 7). These results are not surprising either. The extremely high incidence of missing values in the dataset makes predictive MRCAs prone to be relatively old (Fig. 1), which has a proven negative impact on prediction accuracy^4,8^. As such, simulations show that only species whose predictive MRCA is younger than ∼10% of the height of the phylogeny are expected to show consistently accurate predictions (i.e. at least 75% of *P*^*2*^_sim_ values ≥ 0.75) under a scenario of ‘strong’ phylogenetic signal (i.e. λ = 1)^4^.

It is very exciting to see a burgeoning interest in connecting phylogenetic information with human well-being, but researchers should be clear on the limitations of phylogenetic predictive methods and utilize imputed information with caution and restraint, especially if the goal is employing individual predictions separately (e.g. to evaluate if a species has potential to be ‘source’ of a B vitamin, as in Cantwell-Jones et al.^1^). The dataset gathered by Cantwell-Jones et al.^1^ is impressive, and it would not be surprising if the 1044 species they proposed as candidate sources attract the attention of professionals willing to invest resources in measuring their B-vitamin content.

Unfortunately, the reliability of the predictions that informed the list of candidate species ranges between weak to very weak. While the authors did not intend to predict the exact nutrient content of species but to evaluate if the predicted values were above or below a certain threshold, the fact that most of their predictions are indistinguishable from pure randomness proves the weakness of the inference drawn from imputed information in their study. We hope this commentary serves as a cautionary note for future phylogenetic imputation exercises to carefully assess whether the data meet the requirements for the predictions to be valuable, or at least more accurate than expected by chance.

## Supporting information

Supplementary Software 1-5

## Acknowledgements

We thank the Scientific Computation Centre of Andalusia (CICA) for the computing services they provided. I.M.-C. acknowledges funding from the Spanish Ministry for Science and Innovation (grant no. PID2019-109711RJ-I00 to I.M.-C.) as well as from Comunidad de Madrid and Universidad de Alcalá (funders of I.F.W. through grant CM/BG/2021-003 to I.M.-C.).

## Author contributions

R.M.-V. conceived and discussed the idea with M.A.R. and I.M.-C., performed the calculations and led the writing, I.M.-C. drew the figure, and all the authors contributed to the writing.

## Additional information

### Supplementary information

The online version contains supplementary material available at XXX

## Data availability

All the datasets generated for this study are available from the Figshare Digital Repository at https://doi.org/10.6084/m9.figshare.19682880.v1.

**Correspondence** should be addressed to R.M.-V.

