## Supplementary Information for "Significant phylogenetic signal is not enough to trust phylogenetic predictions"

Supplementary Methods: **Assessing prediction accuracy across candidate species**. We used the method described in Molina-Venegas et al.^4^ to evaluate the expected accuracy of the individual predictions that informed the list of candidate species proposed in Cantwell-Jones et al.^1^ as promising key sources of B vitamins. Given a phylogeny and a continuous trait (concentration of a given B vitamin per species) with missing values (species for which the concentration of the given vitamin is unknown), a Pagel’s λ model of evolution is fitted to the observed data, and the model parameters —i.e. fitted value of lambda (λ_obs_), instantaneous variance of the process (sig^2^), and ancestral state at the root node (z_0_)— are retained. Then, the off-diagonal elements of the phylogenetic variance–covariance matrix are downweighed by λ_obs_, and the so-rescaled phylogeny (which includes both nutritionally known and unknown species) is used to simulate a high number of traits (e.g. n = 1000) using a Brownian motion model of evolution at the estimated sig^2^ and z_0_ parameters. This procedure simulates traits that, on average, show the same degree of phylogenetic signal (hereafter ‘λ_sim_’) as the observed values of the real trait (λ_obs_). However, λ_sim_ scores above or below λ_obs_ are expected due to the stochastic nature of phylogenetic random walks. Thus, only traits for which λ_sim_ = λ_obs_ ± 0.025 and significant (*p* ≤ 0.05) in a likelihood-ratio test are retained, and the simulations are conducted iteratively until all the traits meet the conditions. The permitted variation in λ_sim_ scores around λ_obs_ is set arbitrarily by the user (here ±0.025), and such restriction is simply aimed to avoid traits whose phylogenetic signal λ_sim_ differs greatly from the expectation (λ_obs_) due to occasional random drifts. Thus, the lower the permitted variation the better, but note that highly restrictive settings may increase computation time towards prohibitive levels as the probability for the traits to meet the conditions decreases. For each simulated trait, the values corresponding to the missing ones in the real trait are dropped and subsequently predicted with the maximum likelihood ancestral state method (e.g. as in Cantwell-Jones et al.^1^) using the remaining values. Finally, the accuracy of each individual prediction *i* is measured using the *P^2^_sim_* prediction coefficient:

*P^2^_sim_* = 1 - $\frac{{( - )}^{2}}{}$

where and are respectively the predicted and observed simulated value for species *i* and is the sample variance of the observed values of the simulated trait. The distribution of *P^2^_sim_* values of each species can be summarized to obtain an estimate of the expected accuracy of species-level predictions on the unknown values of the real trait. For example, one may trust the vitamin content predicted for a given species if most of the *P^2^_sim_* scores obtained for that species at the fitted parameters (λ_obs_, sig^2^, z_0_) are overall high (e.g. at least 75% of *P^2^*_sim_ values ≥ 0.75). Otherwise, the prediction may be valueless (see Molina-Venegas et al.^4^ for an extended discussion).

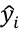

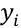

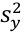

Here, we followed the procedure described above to simulate traits at the parameters estimated for each vitamin (n = 1000 traits per vitamin) and compute a distribution of *P^2^_sim_* values for each vitamin and candidate species (as retrieved from the Supplementary Data 3 in Cantwell-Jones et al.^1^), totaling 1616 distributions (n = 1000 *P^2^_sim_* values per distribution). Then, we repeated the procedure but switching λ_obs_ to zero so that a second set of traits was simulated under a ‘white noise’ model of evolution, which corresponds to a Pagel’s λ model where λ = 0 (absence of any phylogenetic correlation structure)^10^. To ensure complete lack of phylogenetic signal in the ‘white noise’ set, we only retained those traits for which λ_sim_ < 0.001 and *p* > 0.05 in the likelihood-ratio test, and simulations were conducted iteratively until all the traits met the conditions. These traits were used to compute a null distribution of *P^2^_sim_* values (hereafter ‘*P^2^_sim-null_*’) for each vitamin and candidate species. Then, we tested the hypothesis that *P^2^_sim_* values are higher than *P^2^_sim-null_* using two-sample Wilcoxon tests (one test for each B vitamin and candidate species, totaling 1616 tests). Multiple statistical tests may lead to high chance of type I error, and a common procedure to avoid false positives is adjusting *p*-values using Bonferroni correction (e.g. as in Cantwell-Jones et al.^1^). However, this adjustment is conducted at the expense of increasing false negatives^11^. Thus, we presented the statistical evidence using four different scenarios ranging from less conservative to more conservative: (i) exact *p*-values and 5% nominal alpha, (ii) exact *p*-values and 0.1% nominal alpha, (iii) adjusted *p*-values and 5% nominal alpha, and (iv) adjusted *p*-values and 0.1% nominal alpha (Supplementary Data 6). All the analyses were conducted in R^12^ using the packages ‘phytools’^13^, ‘geiger’^14^, and ‘Rphylopars’^15^, and the R code to replicate the analyses is provided in the Supplementary Material attached online to this article. We used Rphylopars to conduct phylogenetic predictions instead of the method employed by Cantwell-Jones et al.^1^ because the former is a reference software for phylogenetic imputation^8,16,17^ and performs notably faster than the latter. Both methods are based in maximum likelihood ancestral state reconstruction and thus produce almost identical predictions (see Supplementary Software 5 for a demonstration).
