## Supplementary Software 1-5 for "Significant phylogenetic signal is not enough to trust phylogenetic predictions": Sup_Software_1_Script_to_randomize_traits.txt

rm(list=ls())

require(phytools)
require(geiger)
fullTree <- read.tree(".../Tree.txt") # This is phylogeny of edible species
fullDat <- matrix(NA,ncol = 500, nrow = 280)

load(".../nutDat.RData") # This is the original dataset with unnecessary columns already removed. It includes 280 rows (species) and 6 columns (vitamins). Contains NAs.

set.seed(NULL)

sort(row.names(nutDat))->sorting

nutDat <- nutDat[sorting,]

row.names(fullDat) <- row.names(nutDat)
colnames(fullDat)  <- paste("run",1:500,sep="")


#Dat_ori <- nutDat[,1] # Remove one "#" at a time to unlock. This will index observed values for thiamine
#Dat_ori <- nutDat[,2] # Remove one "#" at a time to unlock. This will index observed values for riboflavine
#Dat_ori <- nutDat[,3] # Remove one "#" at a time to unlock. This will index observed values for niacin
#Dat_ori <- nutDat[,4] # Remove one "#" at a time to unlock. This will index observed values for panthotenic acid
#Dat_ori <- nutDat[,6] # Remove one "#" at a time to unlock. This will index observed values for folate

names(Dat_ori) <- row.names(nutDat)

which(is.na(Dat_ori)) -> NA_pos

Dat_ori_no_NA <- Dat_ori[!is.na(Dat_ori)]

Tree <- drop.tip(fullTree,fullTree$tip.label[fullTree$tip.label %in% names(Dat_ori_no_NA) == FALSE])

Counter <- 1

while(Counter != 501) {

null_dat <- Dat_ori_no_NA
null_dat <- sample(null_dat)
names(null_dat) <- names(Dat_ori_no_NA)
null_dat <- null_dat[Tree$tip.label]

phylosig(Tree, null_dat, method="lambda", test=TRUE) -> test.i
if(test.i$lambda < 0.001 & test.i$P > 0.05) {

NA_pos[] <- NA

null_dat <- c(null_dat,NA_pos)
null_dat <- null_dat[sorting]

null_dat -> fullDat[,Counter]
Counter <- Counter + 1
print(paste("Trait ", Counter-1," completed",sep=""))
} else
{1}
}

setwd("...") # set working directory to save the data

# Unlock the corresponding line to save the random traits:

#save(fullDat, file="sim_thiamine.RData")
#save(fullDat, file="sim_riboflavin.RData")
#save(fullDat, file="sim_niacin.RData")
#save(fullDat, file="sim_pantothenic.RData")
#save(fullDat, file="sim_folate.RData")
