## Supplementary Software 1-5 for "Significant phylogenetic signal is not enough to trust phylogenetic predictions": Sup_Software_2_Script_to_analyze_randomized_traits.txt

# Script to conduct leave-one-species-out crossvalidation trials on randomized sets and compute 95% CI around predictions. The script is based on the code provided in Cantwell-Jones et al. 2022 #

rm(list=ls())

require(phytools)
require(geiger)

source("/.../phyloPred.jknife.R")

load(".../nutDat.RData") # This is the original dataset with unnecessary columns already removed. It includes 280 rows (species) and 6 columns (vitamins). Contains NAs.

to_sort <- row.names(nutDat) # this will be used to sort the rows of the resultant dataset "pred"

rm(nutDat)

fullTree <- read.tree("/.../Tree.txt") # load the phylogeny

#load("/.../sim_thiamine.RData") # Unlock to analize thiamine. The "sim" dataset was generated using the code "Script_to_randomize_traits.txt"
#load("/.../sim_riboflavin.RData") # Unlock to analize riboflavine. The "sim" dataset was generated using the code "Script_to_randomize_traits.txt"
#load("/.../sim_niacin.RData") # Unlock to analize niacin. The "sim" dataset was generated using the code "Script_to_randomize_traits.txt"
#load("/.../sim_pantothenic.RData") # Unlock to analize pantothenic acid. The "sim" dataset was generated using the code "Script_to_randomize_traits.txt"
#load("/.../sim_folate.RData") # Unlock to analize folate. The "sim" dataset was generated using the code "Script_to_randomize_traits.txt"

datSp <- rownames(fullDat)

exclude <- fullTree$tip.label[which(match(fullTree$tip.label, datSp, nomatch=0)==0)]

tree <- drop.tip(fullTree, exclude) 

nutDat<-fullDat

nut<-data.frame(sapply(colnames(nutDat), function(x) log(nutDat[,x]+0.01)))

rownames(nut)<-rownames(nutDat)

nutList<-lapply(1:ncol(nut), function(x) nut[which(is.na(nut[,x])==FALSE),x])
	names(nutList)<-colnames(nut)
	for(i in 1:length(nutList)){names(nutList[[i]])<-rownames(nut)[which(is.na(nut[,i])==FALSE)]}
	
nutList.n<-sapply(1:length(nutList), function(x) length(nutList[[x]]))

treeList<-lapply(1:length(nutList), function(x) drop.tip(tree, which(match(tree$tip.label, names(nutList[[x]]), nomatch=0)==0)))

errorMat<-matrix(,length(nutList),3, dimnames=list(names(nutList), c("median_diff_percMean", "median_diff_nSD", "errorRate")))

setwd("/.../Temp") # set working directory to save the results

for(i in 1:500) {  # Note that this loop will take a very long time, and thus using a supercomputation cluster is highly recommended.

	nut<-nutList[[i]]
	names(nut)<-names(nutList[[i]])		

	pred<-matrix(, length(nutList[[i]]), 6, dimnames=list(names(nutList[[i]]), c("measuredVal", "predVal", "lcl_predInt", "ucl_predInt", "diff_percMean", "diff_nSampleSD")))
	withinCIMat<-matrix(, length(nutList[[i]]), 1, dimnames=list(names(nutList[[i]]), "withinCI"))
		
	for (j in 1:length(nutList[[i]])){

		result<-phyloPred.jknife(treeList[[i]], nut, names(nut)[j])

		predVal<-as.numeric(result$PredVal)
		measVal<-as.numeric(nut[j])
		diff<-abs(measVal - predVal)
		predSE<-as.numeric(result$PredSE)
		diff_nPredSE<-diff / predSE
		if (diff_nPredSE < 2) withinCIMat[j]<-TRUE else withinCIMat[j]<-FALSE
		
		lclPredInt<-predVal - (1.96 * predSE)
		uclPredInt<-predVal + (1.96 * predSE)	
		
		predVal<-exp(predVal) - 0.01
		measVal<-exp(nut[j]) - 0.01
		lclPredInt<-exp(lclPredInt) - 0.01
		uclPredInt<-exp(uclPredInt) - 0.01
		nutSD<-sd(exp(nut) -0.01)

		if (measVal == 0) diff1<-NA else diff1<-as.numeric((abs(measVal - predVal) / measVal) * 100)
		if (measVal == 0) diff2<-NA else diff2<-as.numeric((abs(measVal - predVal) / nutSD))
						
		pred[j,]<-round(c(measVal, predVal, lclPredInt, uclPredInt, diff1, diff2), 3)

	}

	pred<-data.frame(pred, withinCIMat)
	
	pred <- pred[,c(1:4,7)]
	
    pred <- pred[to_sort,]
	
save(pred, file=paste("Res_",i,".RData",sep="")) 

print(paste("Dataset ",i," completed", sep=""))

}
