## Supplementary Software 1-5 for "Significant phylogenetic signal is not enough to trust phylogenetic predictions": Sup_Software_3_Script_method_Molina_Venegas_2018.txt

rm(list=ls())

require(Rphylopars)
require(phytools)
require(geiger)

fullTree <- read.tree("/.../Tree.txt") # This is the phylogeny of edible species

fullDat <- read.csv("/.../B_vitamin_nutrition_data.csv")  # This is the original dataset

#proposed <- read.table("/.../Proposed_Thiamine.txt",row.names=1,dec=",",header=TRUE) # This is the list of species representing promising key sources of thiamine as suggested by the authors (Supplementary Data 3 in Cantwell-Jones et al. 2022). Unlock this line to analyze thiamine   
#proposed <- read.table("/.../Proposed_Riboflavin.txt",row.names=1,dec=",",header=TRUE) # This is the list of species representing promising key sources of riboflavin as suggested by the authors (Supplementary Data 3 in Cantwell-Jones et al. 2022). Unlock this line to analyze riboflavin   
#proposed <- read.table("/.../Proposed_Niacin.txt",row.names=1,dec=",",header=TRUE) # This is the list of species representing promising key sources of niacin as suggested by the authors (Supplementary Data 3 in Cantwell-Jones et al. 2022). Unlock this line to analyze niacin  
#proposed <- read.table("/.../Proposed_Folate.txt",row.names=1,dec=",",header=TRUE) # This is the list of species representing promising key sources of folate as suggested by the authors (Supplementary Data 3 in Cantwell-Jones et al. 2022). Unlock this line to analyze folate  

rownames(fullDat) <- fullDat[ , 1] #make species names rownames of nutrition data

fullDat <- fullDat[ , -c(1:3, 5)] #remove unnecessary columns 

#vitamin <- fullDat[,1] # column 1 for thiamine. Unlock this line to analyze thiamine
#vitamin <- fullDat[,2] # column 1 for riboflavin. Unlock this line to analyze riboflavin
#vitamin <- fullDat[,3] # column 1 for niacin. Unlock this line to analyze niacin
#vitamin <- fullDat[,6] # column 6 for folate. Unlock this line to analyze folate

names(vitamin) <- row.names(fullDat)

vitamin_val <- na.omit(vitamin)

##### To obtain the parameters of the lambda model fitted to the observed values of the trait #####

Tree_vit <- drop.tip(fullTree,fullTree$tip.label[fullTree$tip.label %in% names(vitamin_val) == FALSE])
 
vitamin_val_par <- vitamin_val[Tree_vit$tip.label]

lambdaFit_vit <- fitContinuous(Tree_vit, vitamin_val_par, model="lambda")$opt

lambda <- lambdaFit_vit$lambda # phylogenetic signal

sig2 <- lambdaFit_vit$sigsq # instantaneous variance of the process

z0 <- lambdaFit_vit$z0 # ancestral state at the root node

####################################################################################################

vitamin <- c(vitamin_val,rep(NA,dim(proposed)[[1]]))

names(vitamin)[(length(vitamin_val)+1):length(vitamin)] <- row.names(proposed)

Tree <- fullTree

Tree <- drop.tip(Tree, Tree$tip.label[Tree$tip.label %in% names(vitamin) == FALSE])

Trait <- vitamin

Trait <- Trait[Tree$tip.label]

range.lambda = 0.025

set.seed(NULL)

for(i in 1:1000) { # This loop may take a very long time. Using a supercomputation cluster is strongly recommended

km <- 0

while(km == 0){

x <- fastBM(rescale(Tree,"lambda", lambda), a=z0, sig2=sig2, bounds=c(0,Inf)) # simulates trait using the parameters of the lambda model (bound to a minimum value of zero because negative concentrations of B vitamins are not possible) 

Signal <- phylosig(Tree, x, "lambda", test=TRUE) # computes phylogenetic signal (Pagel's lambda) and likelihood ratio test on the simulated trait

if(Signal$lambda >= (lambda - range.lambda) & Signal$lambda <= (lambda + range.lambda) & Signal$P <= 0.05) {km <- 1} else # conditions to retain trait

{1}

}

trait <- data.frame(Tree$tip.label,x[names(x)%in%Tree$tip.label],x[names(x)%in%Tree$tip.label]) # phylopars requires a minimum of two traits in the input, thus it is duplicated in the dataframe "trait"    

colnames(trait)<-c("species","V1","V2")

trait[row.names(trait)[row.names(trait)%in%names(Trait[is.na(Trait)])],2:3] <- NA # this values are missing in the real dataset 

PE <- phylopars(trait, Tree, model = "lambda", pheno_error=FALSE, pheno_correlated = FALSE, phylo_correlated=FALSE) # runs phylopars using only the phylogeny and a lambda model 

PE_predict <- PE$anc_recon

Predict_pars <- PE_predict[1:length(Tree$tip.label),1]

x_miss <- x[names(Trait[is.na(Trait)])]

Predict_pars_miss <- Predict_pars[names(Trait[is.na(Trait)])]

DF <- data.frame(x_miss,Predict_pars_miss)

P2 <- 1 - (((DF[,2] - DF[,1])^2) / var(x)) # prediction coefficient for each species 

Result <- list(sim_observed = x_miss, sim_predicted = Predict_pars_miss, accuracy_val = P2, trait_sim = x, Signal_x = Signal)

setwd("/.../") # set working directory to save the results

save(Result, file = paste("Result_",i,".RData",sep=""))

print(paste("Loop ",i," completed", sep=""))

}
